# Evaluation of haplotype-aware long-read error correction with hifieval

**DOI:** 10.1101/2023.06.05.543788

**Authors:** Yujie Guo, Xiaowen Feng, Heng Li

**Affiliations:** Department of Data Science, Dana-Farber Cancer Institute, Boston, MA, USA, 02215; Department of Biomedical Informatics, Harvard Medical School, Boston, MA, USA, 02215

## Abstract

**Summary:** The PacBio High-Fidelity (HiFi) sequencing technology produces long reads of *>*99% in accuracy. It has enabled the development of a new generation of *de novo* sequence assemblers, which all have sequencing error correction as the first step. As HiFi is a new data type, this critical step has not been evaluated before. Here, we introduced hifieval, a new command-line tool for measuring over- and under-corrections produced by error correction algorithms. We assessed the accuracy of the error correction components of existing HiFi assemblers on the CHM13 and the HG002 datasets and further investigated the performance of error correction methods in challenging regions such as homopolymer regions, centromeric regions, and segmental duplications. Hifieval will help HiFi assemblers to improve error correction and assembly quality in the long run.

**Availability and implementation:** The source code is available at https://github.com/magspho/hifieval

**Supplementary information:** Supplementary data are available at *Bioinformatics* online.

## 1. Introduction

The PacBio High-Fidelity (HiFi) sequencing technology produces long reads of *∼*15 kilobases (kb) in length and *>*99% in accuracy (Wenger *et al*., 2019). This new data type leads to the development of several recent sequence assemblers, including HiCanu (Nurk *et al*., 2020), hifiasm (Cheng *et al*., 2021, 2022), mdBG (Ekim *et al*., 2021), LJA (Bankevich *et al*., 2022), and Verkko (Rautiainen *et al*., 2023; Rautiainen and Marschall, 2021), which leverage the high accuracy of HiFi reads and could generate phased assemblies of higher quality than previous methods (Jarvis *et al*., 2022). HiFi has also played a central role in the complete assembly of the first human genome (Nurk *et al*., 2022; Rhie *et al*., 2023).

The power of HiFi read assembly mainly comes from resolving similar repeats or haplotypes by requiring exact sequence matches between reads (Nurk *et al*., 2020; Cheng *et al*., 2021). Although HiFi reads are accurate, there are still a small number of sequencing errors per read. These errors would result in inexact matches and fail the assembly algorithms. To obtain near error-free reads, all the HiFi assemblers mentioned above start with error correction (EC), a critical step to correct away most sequencing errors in reads. Errors at the EC step may be propagated to the assembly step and lead to broken contigs or misassemblies downstream.

Zhang *et al*. (2020) evaluated many EC tools developed for correcting noisy long reads. However, most of these tools disregard phasing and would collapse reads originated from different repeat copies or from different parental haplotypes in a diploid sample. They are not used by modern assemblers and are largely obsolete. To better understand the accuracy of modern EC tools, we conducted a new benchmark using real human data. Different from Zhang *et al*. (2020) who did evaluation on small homozygous model organisms, we took high-coverage HiFi reads both from a homozygous cell line CHM13 and from a diploid individual HG002. More importantly, we developed a user-facing tool, hifieval, to measure and record under-corrected and over-corrected bases in reads. This allows other assembler developers to associate EC errors with different genomic features and to study the behavior of EC tools in detail.

## 2. Methods

Suppose we have the perfect assembly of a sample and a set of reads sequenced from the same sample. After EC, we expect corrected reads to be exactly aligned to the assembly. Any substitutions and gaps in the alignment would be EC errors. Hifieval makes use of this observation to evaluate correction errors.

In practice, the assembly may have errors and the sample may contain somatic mutations. We would overestimate errors due to imperfect assembly. Nonetheless, the assembly tends to be more accurate than EC as the assembly is built from multiple sequencing technologies and involves other downstream algorithms such as graph cleaning and consensus which can prune uncorrected errors. We can capture the majority of EC errors even if the assembly has minor errors.

### 2.1. Evaluating tools without homopolymer compression

We align the raw reads and the corrected reads to the ground truth assembly using minimap2 (Li, 2018). For each read, hifieval compares the alignment of the raw read and the alignment of the corrected read and calculates three metrics: correct corrections (CC), errors that are in raw reads but not in corrected reads; under-corrections (UC), errors present in both raw and corrected reads; and over-corrections (OC), new errors found in corrected reads but not in raw reads. These are analogous to true positive (TP), false negative (FN), and false positive (FP), respectively (Fig 1). Therefore, we have the following definitions:

**Fig. 1:**
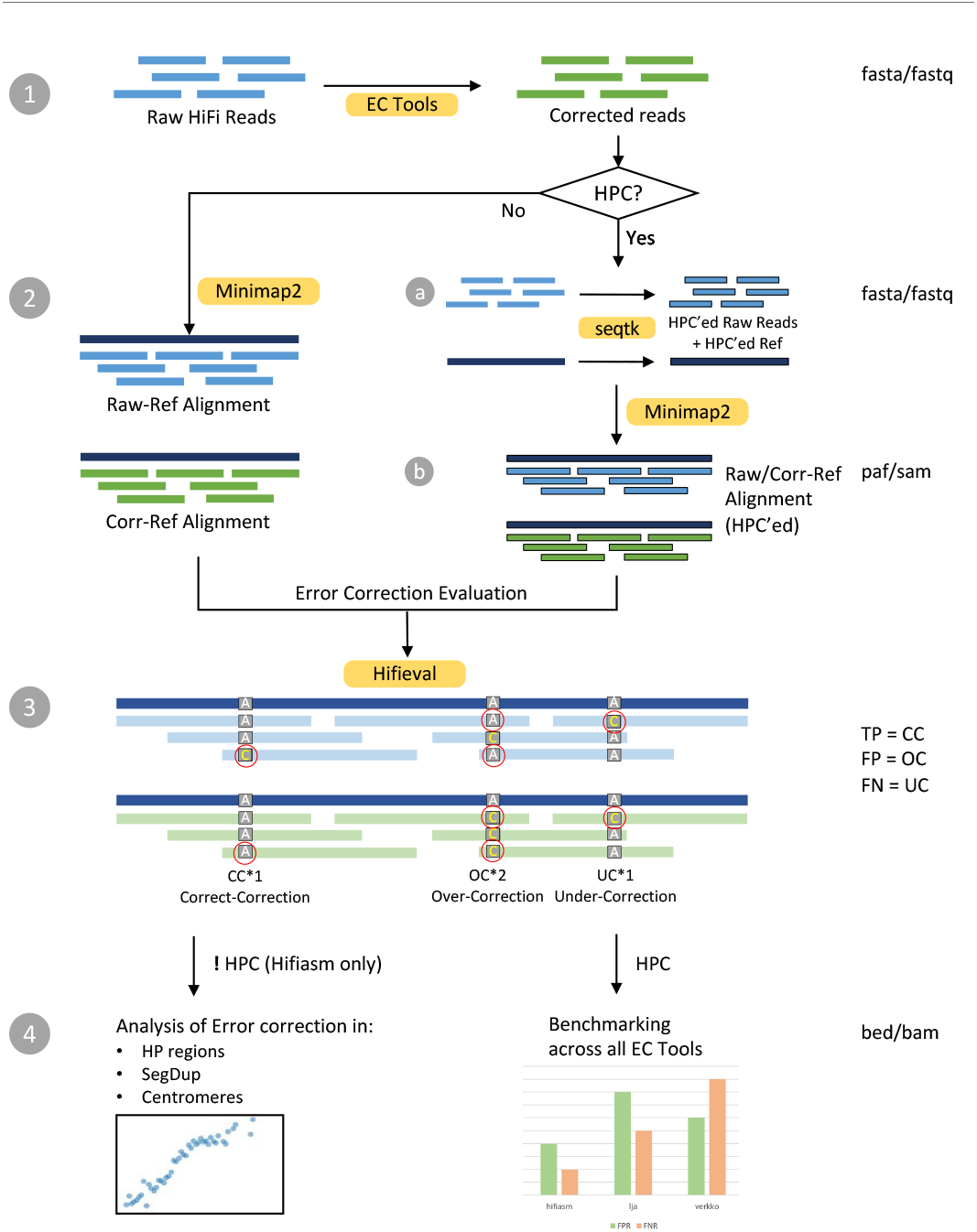
Hifieval workflow.

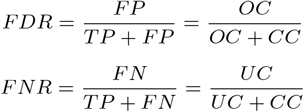

Occasionally, raw reads and corrected reads may be mapped to different positions. Hifieval computes CC, OC and UC anyway but we did not count them in our evaluation.

### 2.2. Evaluating tools with homopolymer compression

Homopolymer compression (HPC) converts a run of identical bases, called a homopolymer, to a single base. It is a simple and widely used method for reducing the negative impacts of homopolymer expansion/contraction error from long-read sequencing technologies. Some EC tools such as Verkko and LJA only report HPC reads. To make the alignment consistent, we also apply HPC to both the truth assembly and the raw reads and perform minimap2 alignment all in the HPC encoding. CC, UC and OC can be measured in a similar manner to the previous section.

## 3. Results

### 3.1. Datasets

#### 3.1.1. Simulated reads

We used *Escherichia coli* str. K-12 substr. MG1655 genome as our reference to simulate the reads (AC:NC 000913.3). We simulated HiFi reads at 30-fold coverage using PBSIM2 with a uniform error rate of 0.2% across the reference (Ono *et al*., 2021). This resulted in 8970 reads from one reference chromosome.

#### 3.1.2. Real reads

We included two real human datasets: T2T-CHM13 and HG002. For CHM13, We took the published assembly (Nurk *et al*., 2022) without the Y chromosome as ground truth and obtained reads from SRR11292120–SRR11292123 at 30-fold coverage. For HG002, we took the Verkko assembly (Rautiainen *et al*., 2023) as the ground truth and downloaded reads from SRR10382244, SRR10382245, SRR10382248 and SRR10382249. These four runs gave 34-fold coverage in total.

### 3.2. Evaluated EC tools

We evaluated three EC tools: hifiasm (Cheng *et al*., 2021), LJA (Bankevich *et al*., 2022), and Verkko (Rautiainen *et al*., 2023). While LJA and Verkko only output corrected reads in the HPC encoding, hifiasm preserved homopolymer runs. For a consistent comparison between the three tools, we emulated hifiasm HPC correction, called hifiasmhpc, by applying hifiasm to homopolymer-compressed raw reads.

HiCanu (Nurk *et al*., 2020) and mdBG (Ekim *et al*., 2021) are also optimized for HiFi assembly and both have an error correction step. We did not evaluate HiCanu because it used the same error correction module as Verkko. mdBG corrected reads in the minimizer space. It did not output in the base space. In theory, EC tools developed for noisy long reads (Zhang *et al*., 2020) could work for accurate PacBio HiFi reads. Nonetheless, most of these older EC tools were developed for small genomes and did not scale well to human data. More importantly, they disregarded phasing and would not be useful for high-quality phased assembly.

### 3.3. Evaluating simulated *E. coli* reads

As a sanity check, we started with simulated *E. coli* dataset. Both hifiasm and LJA performed well. Verkko reported many over-corrections with 25% of its corrections being wrong (Fig. S10). We looked at the results in IGV but could not figure out why it made these many errors.

### 3.4. Evaluating on the CHM13 dataset

For each EC tool, only *∼*0.2% of corrected reads were mapped to positions different from raw reads. These reads contributed to

*∼*0.5% of total number of corrections and thus would have little effect on the total number of OC, UC and CC. We excluded these reads when calculating FNR and FDR.

On this dataset, hifiasm appeared to have lower FNR and FDR than hifiasmhpc (Fig. 2A). This was because hifiasm made nine times as many CC as hifiasmhpc and thus had a much larger denominator when we calculated FNR and FDR. EC tools applying HPC are not directly comparable to EC tools without HPC.

**Fig. 2:**
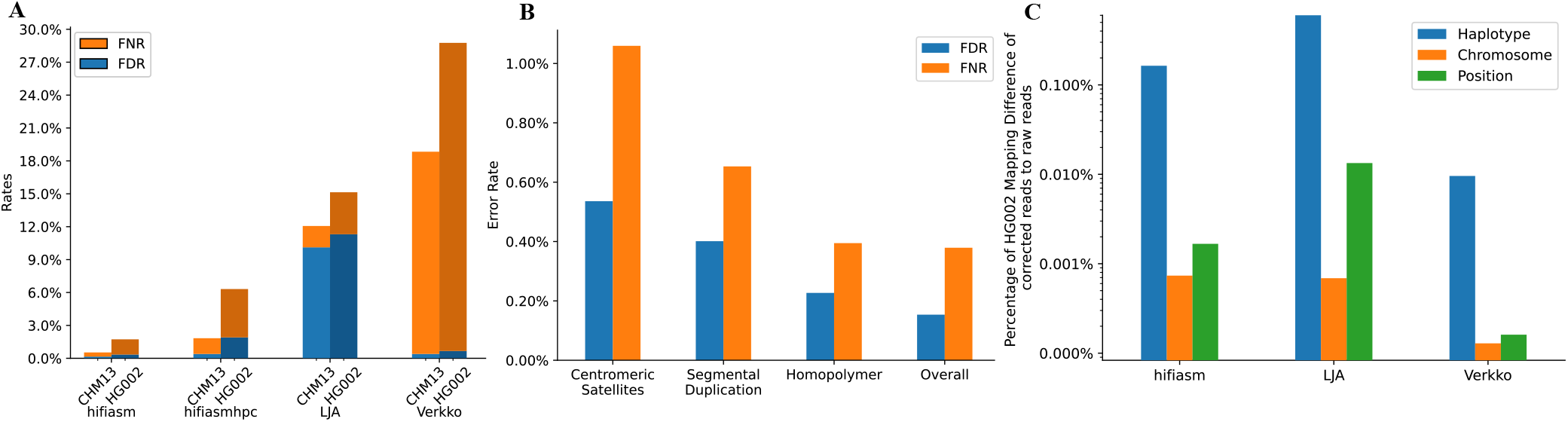
A) The FNR and FDR of Reads Error Correction of each tool on CHM13 and HG002 respectively. B) Hifiasm EC performance in CHM13 challenging regions. C) Percentage of HG002 Mapping Difference of corrected reads to raw reads against Verkko assembled reference. Reads are filtered such that at least either one of raw or corrected read has mapping quality *≥* 2.

Between the three EC tools applying HPC, hifiasmhpc and Verkko had similar FDR at *<*0.4%. LJA had a much higher FDR. It also has more sparse miscorrections (Fig. S1). Verkko on the other hand had a much higher FNR (Fig. S8). 77% of Verkko UCs came from with reads with 10 or more UCs. For these reads, Verkko often corrected part of a read perfectly without touching the errors on the rest of the read. Verkko effectively threw away 18% of data though the errors it corrected were mostly right.

For CHM13, we also stratified hifiasm FNR and FDR by annotation (Fig. 2B, S2). As expected, hifiasm had higher error rate in complex regions such as centromeric satellites and segmental duplications. FNR and FDR were only slightly elevated in homopolymer runs.

### 3.5. Evaluating on the HG002 dataset

All three EC tools had higher error rates on the HG002 dataset than on the CHM13 dataset (Fig. 2A) probably because the two similar parental haplotypes confused the tools. Note that the HG002 assembly were not as polished as the CHM13 assembly. Nonetheless, the contig base accuracy was about two orders of magnitude higher than HiFi base accuracy. Therefore, the great majority read-to-contig differences we saw were sequencing errors. Our evaluation method was still applicable.

For this diploid sample, we counted reads that were mapped to different positions before and after EC and classified the mapping differences into three categories: haplotype difference, chromosome difference and position difference. The correction of a read would lead to a haplotype difference if the raw read and the corrected read were mapped to opposite parental haplotypes, and the alignment of the raw or the alignment of the corrected read had a mapping quality 2 or higher. Similarly, we counted a chromosome difference if the raw read and the corrected read were mapped to different chromosomes on the same haplotype and counted a position difference if the raw read and the corrected were mapped to different positions on the same chromosome.

Fig. 2C showed that Verkko was least likely to correct reads onto the opposite haplotype, at an error rate an order of magnitude lower than hifiasm and LJA (Fig. S4). We checked the haplotype errors made by hifiasm and found that most of the errors occurred in long regions where there were a few heterozygous insertions and deletions (INDELs) but no heterozygous SNPs. These haplotype differences were likely to cause missing INDELs. As we increased the mapping quality threshold to 10, the large haplotype difference between hifiasm and Verkko was much reduced (Fig. S9), suggesting hifiasm and Verkko would rarely correct a read to a very different haplotype. We also performed similar analysis on CHM13 dataset (Fig S5-7).

In Fig. 2A and 2C, we used the Verkko assembly as the ground truth. This could potentially be biased towards Verkko. To evaluate this possibility, we also did the same set of analyses but taking a hifiasm assembly as the ground truth. The results remained similar (Fig. S3). This has confirm the quality of the assembly would not change the overall conclusion.

## 4. Conclusion

Hifieval is a fast tool that performs systematic benchmark of haplotype-aware long read error correction tools. Unlike earlier studies, hifieval evaluates phased assemblies and can distinguish under-corrections and over-corrections. It is perhaps the first user-facing EC evaluation tool that can be easily deployed to users’ own datasets. In our analyses, it is apparent that current EC tools all have their weakness. We hope hifieval can help the development of more accurate EC tools which would be essential to high-quality assembly.

## 6. Author contributions statement

H.L. conceived the project. Y.G. and X.F conducted the experiments. Y.G. analyzed the results. Y.G. and H.L. wrote the manuscript.

## 7. Acknowledgments

This work is supported by US National Institute of Health grant R01HG010040 and U01HG010961 to H.L.

