## Supplementary figures and images for "Evaluation of haplotype-aware long-read error correction with hifieval"

### Fig. S1

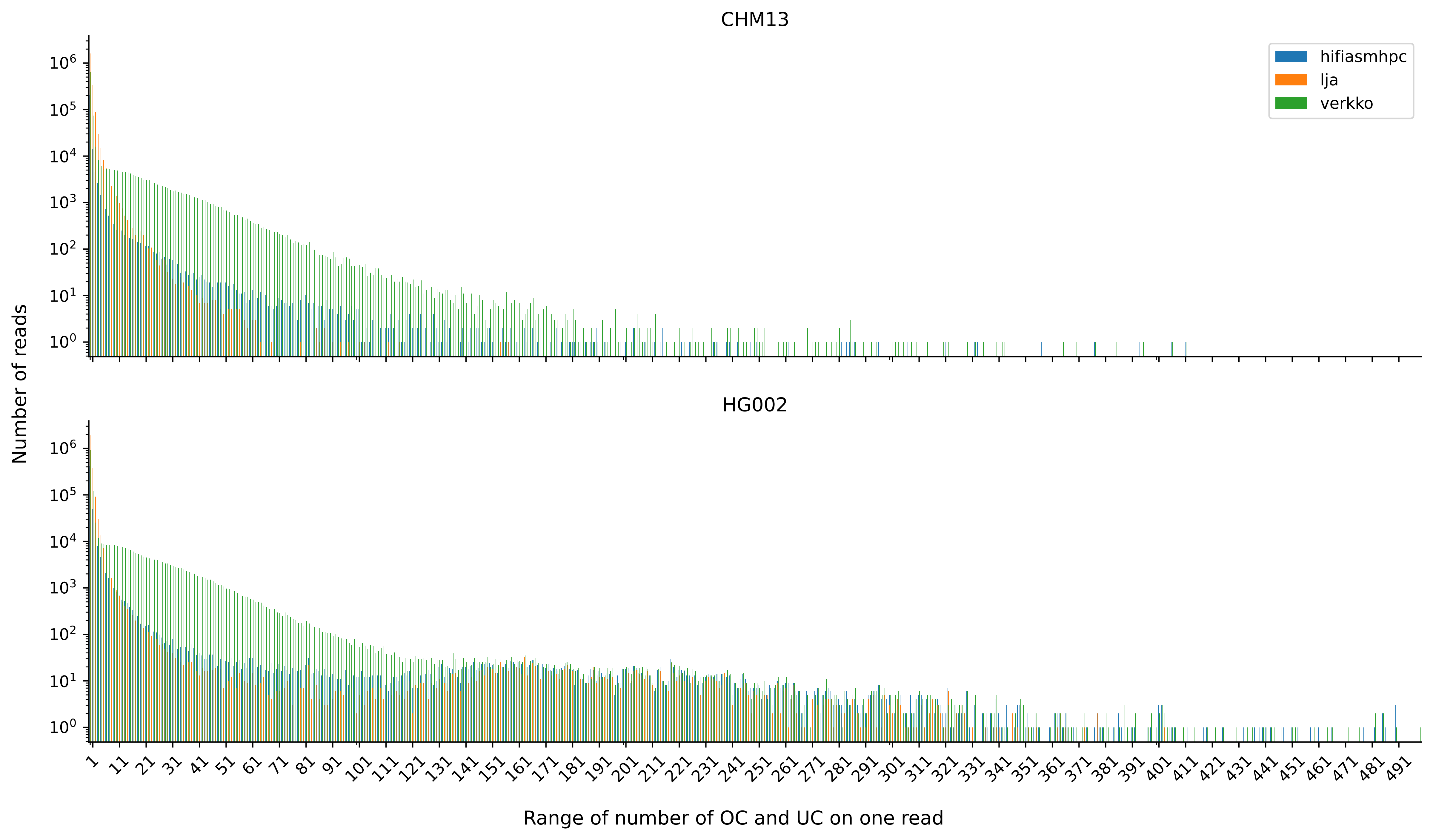

### Fig. S2

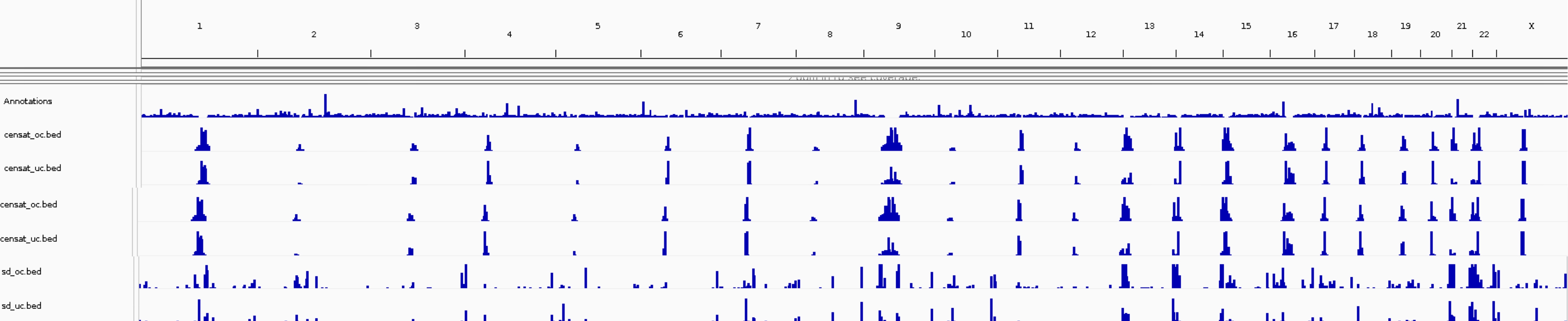

### Fig. S3

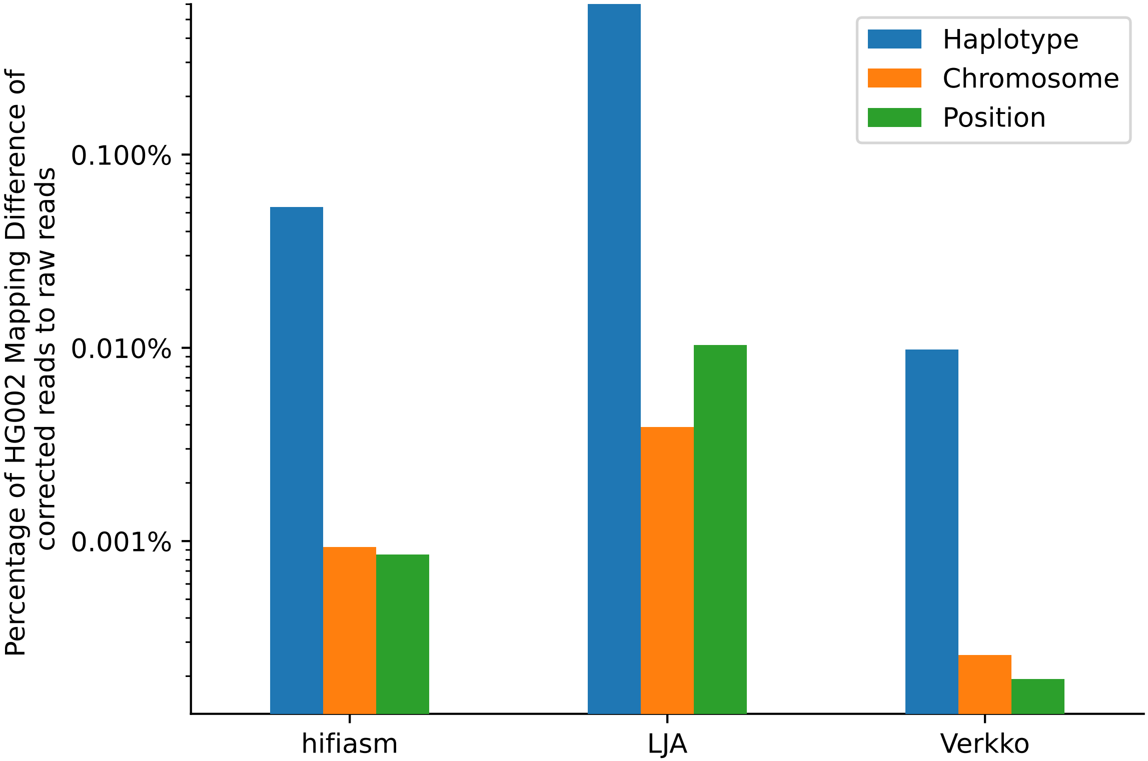

### Fig. S4

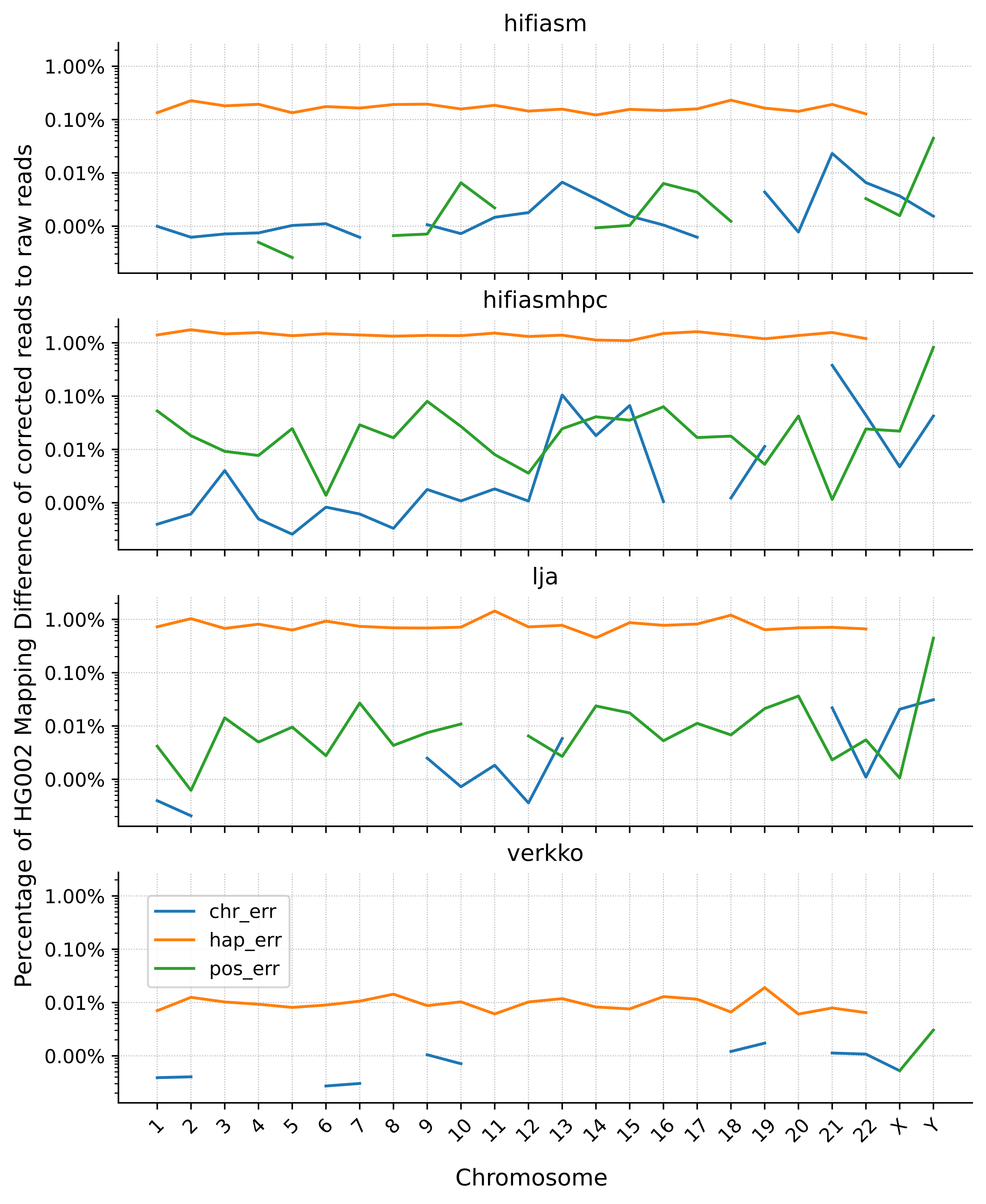

### Fig. S5

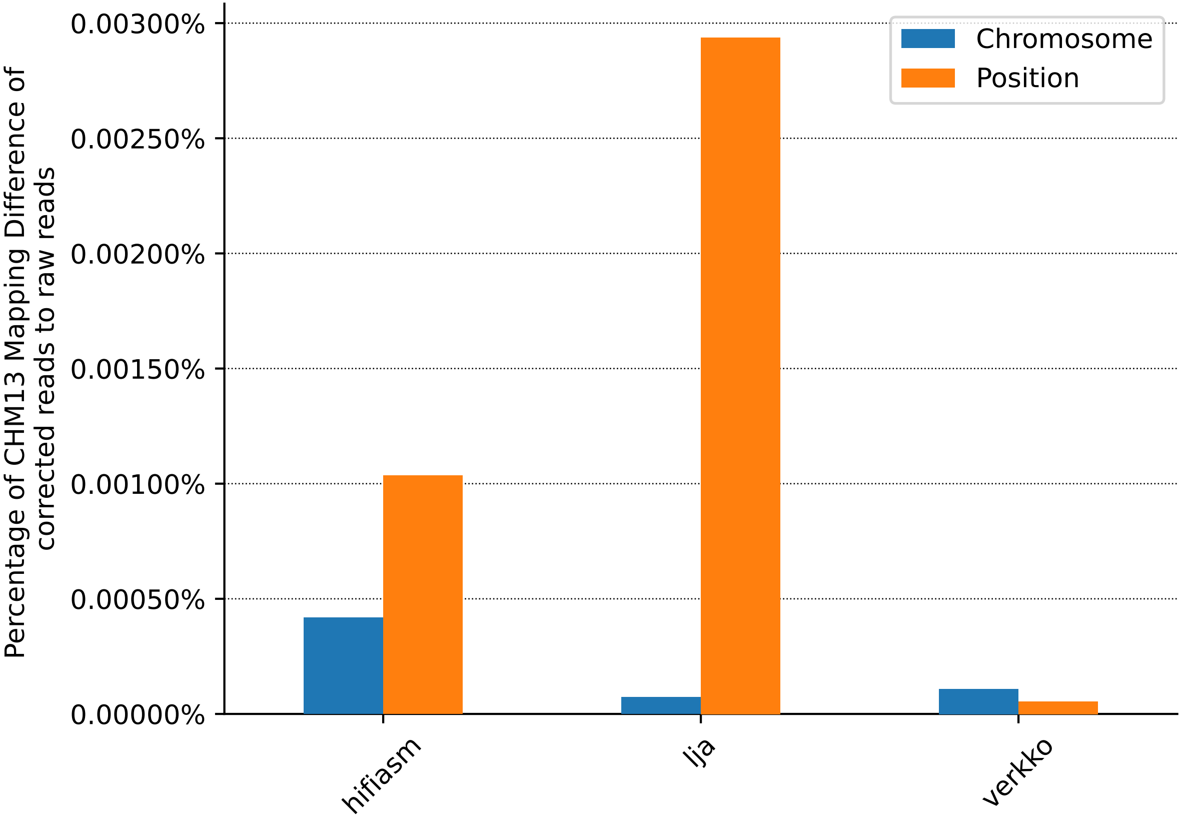

### Fig. S8

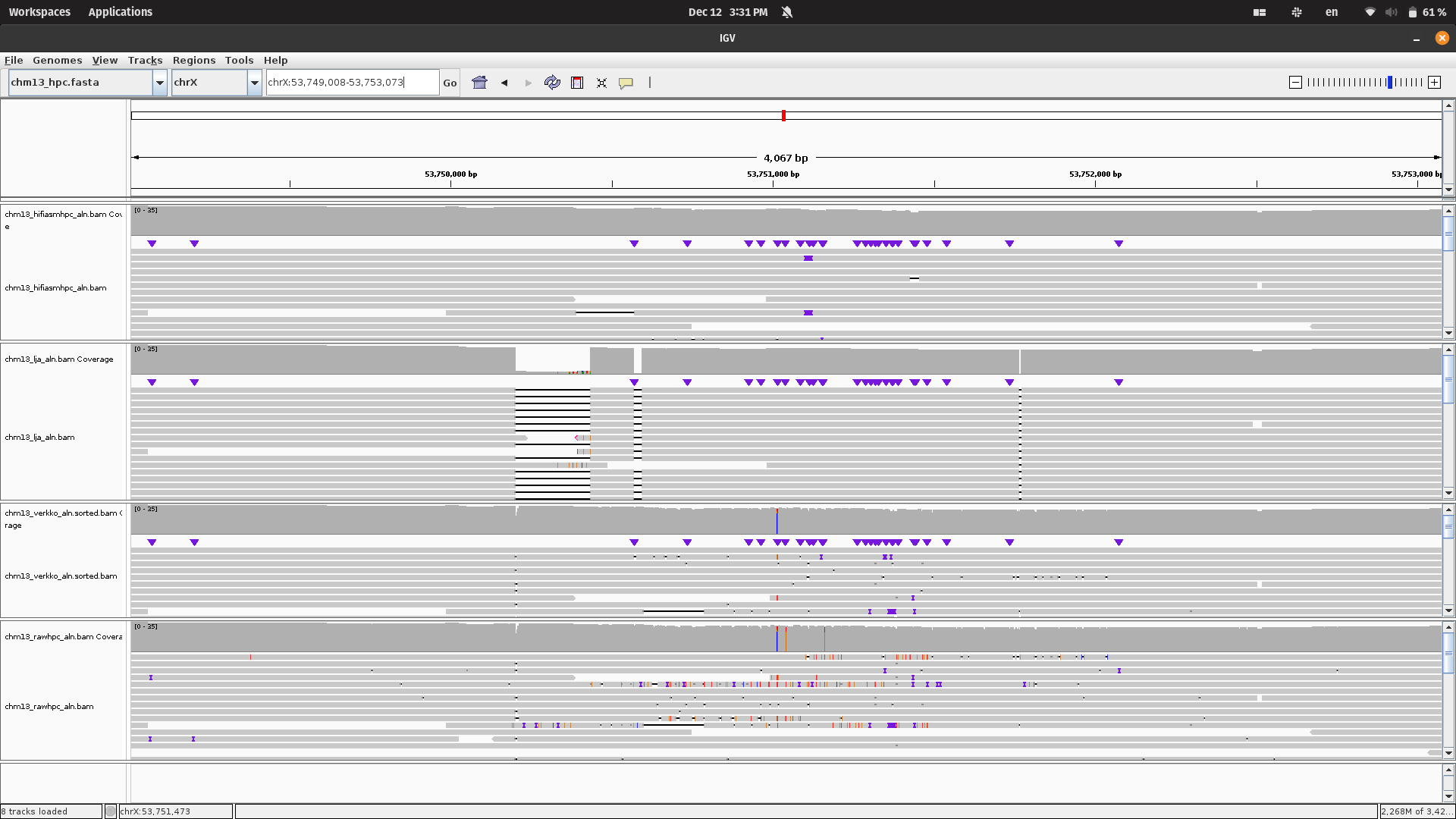

### Fig. S9

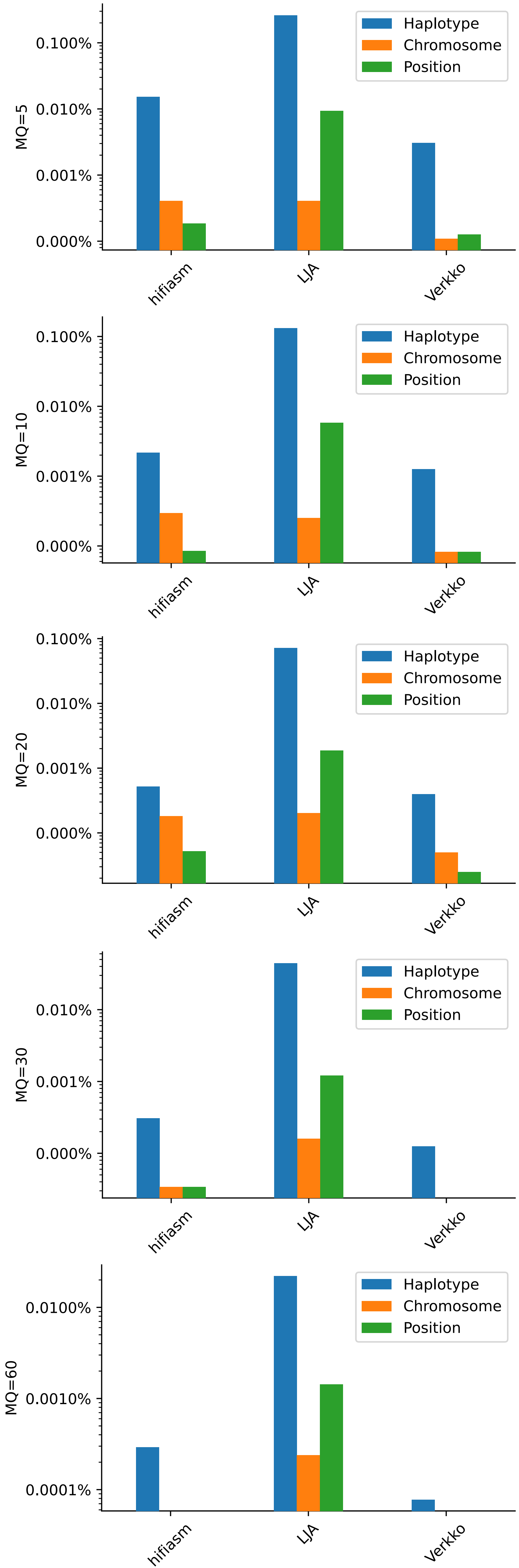

### Fig. S10

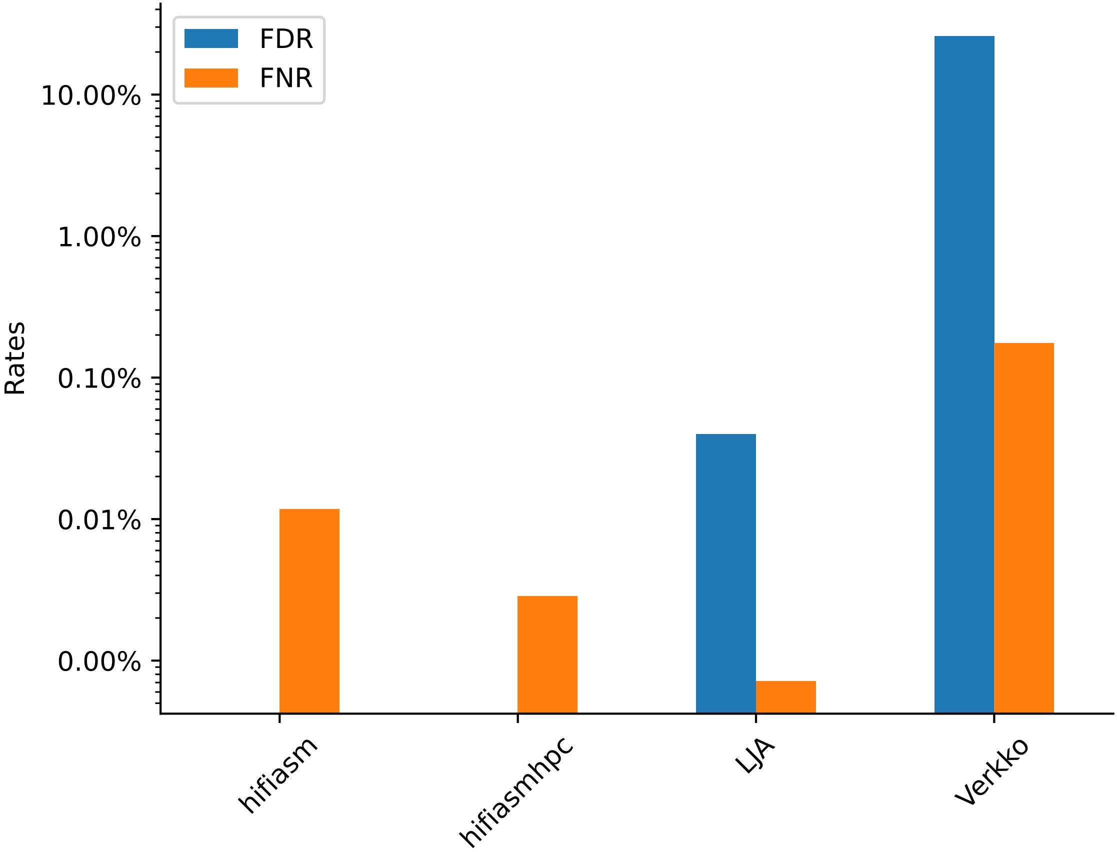
