## Supplementary material for "Evaluation of haplotype-aware long-read error correction with hifieval": Fig. S6

### Mapping Difference between CHM13 raw reads and Hifiasm corrected reads against reference genome

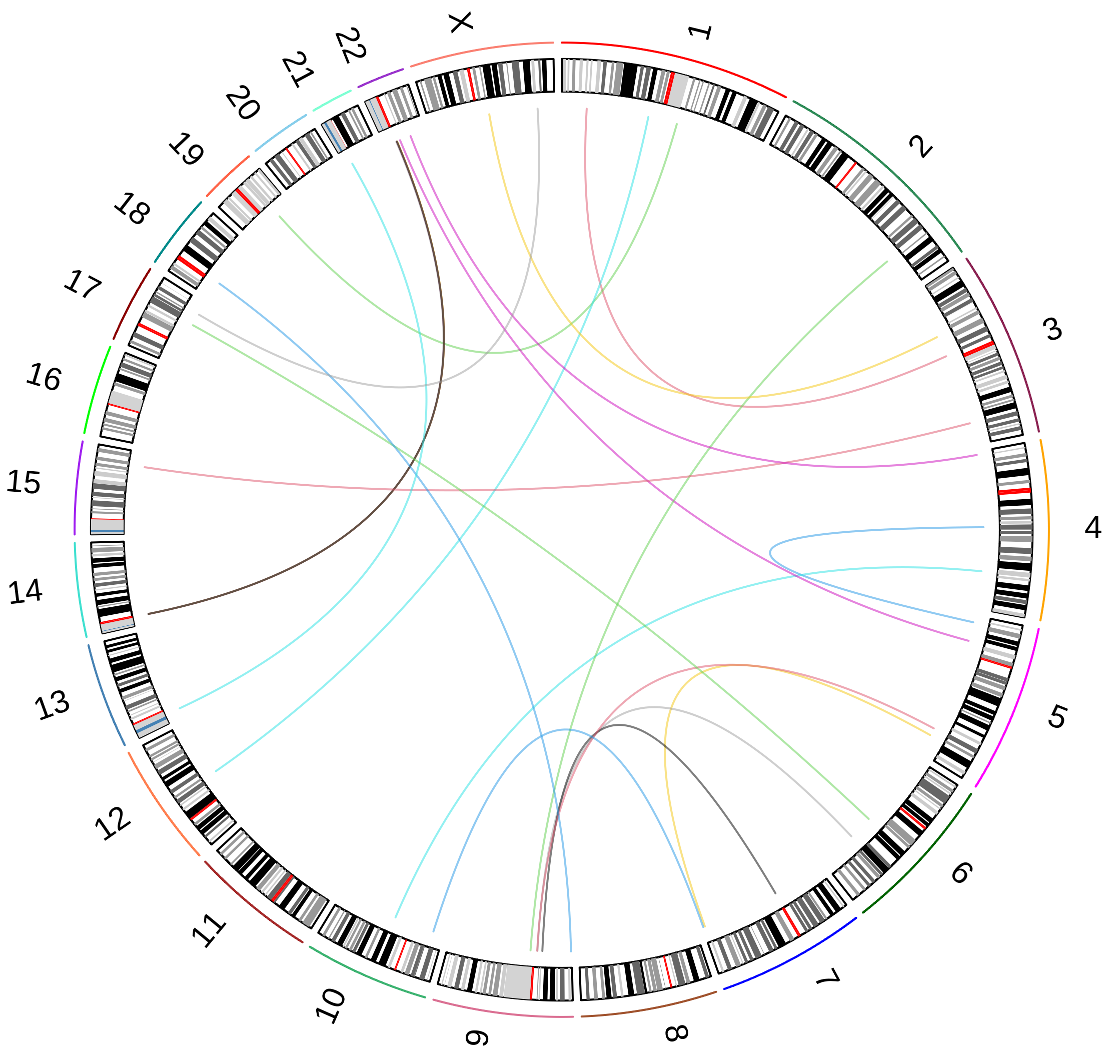
