## Supplementary material for "Evaluation of haplotype-aware long-read error correction with hifieval": Fig. S7

### Mapping Difference between HG002 raw reads and Hifiasm corrected reads against Verkko assembled reference

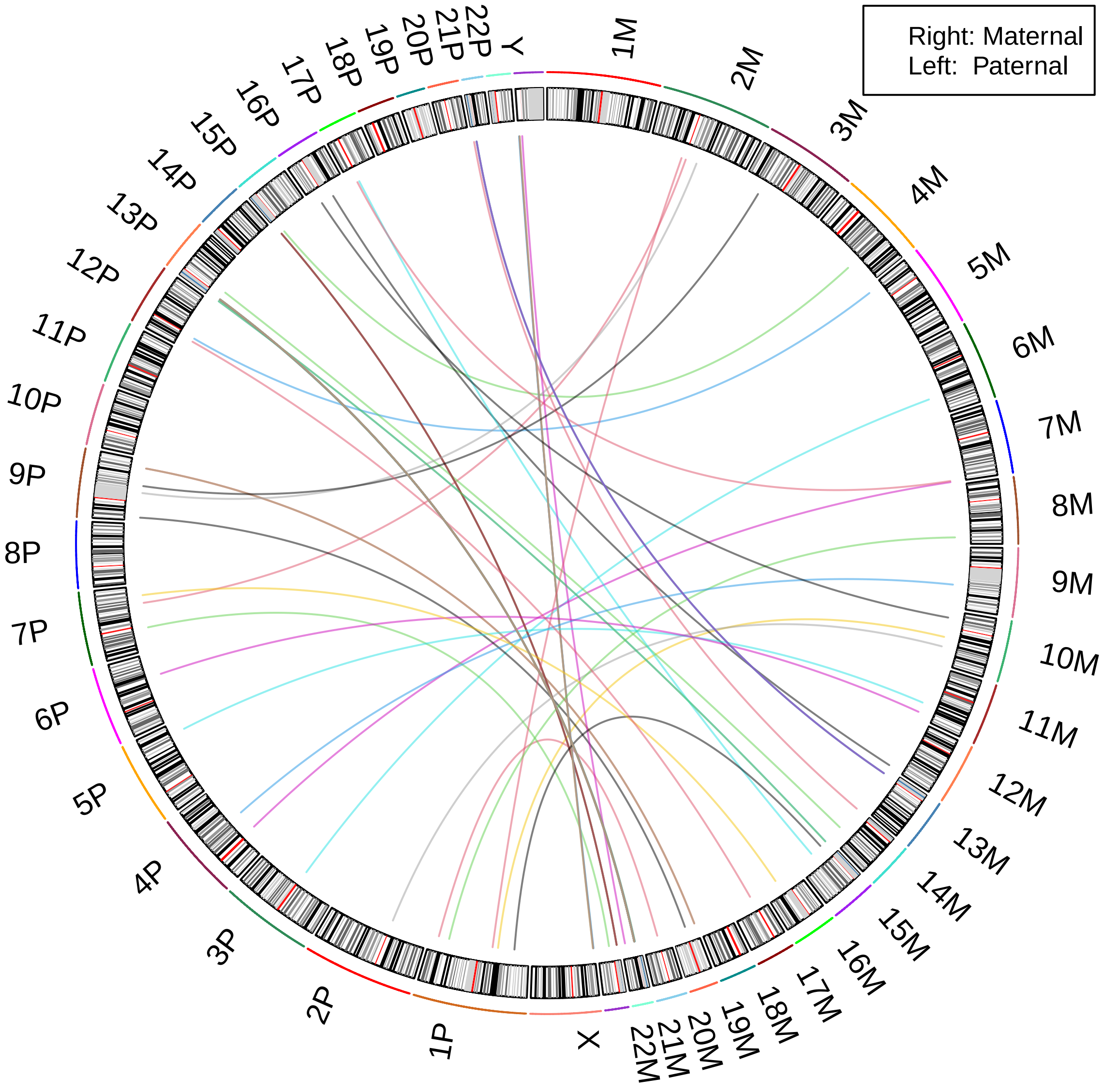
